# Epigenetic change induced by *in utero* dietary challenge provokes phenotypic variability across multiple generations of mice

**DOI:** 10.1101/2020.08.07.241034

**Authors:** Mathew Van de Pette, Antonio Galvao, Steven J. Millership, Wilson To, Andrew Dimond, Chiara Prodani, Grainne McNamara, Ludovica Bruno, Alessandro Sardini, Zoe Webster, James McGinty, Paul French, Anthony G. Uren, Juan Castillo-Fernandez, Rosalind M. John, Anne C. Ferguson-Smith, Matthias Merkenschlager, Gavin Kelsey, Amanda G. Fisher

## Abstract

Transmission of epigenetic information between generations occurs in nematodes, flies and plants, mediated by specialised small RNA pathways, histone H3K9me3, H3K27me3, H4K16ac and DNA methylation^1-3^. In higher vertebrates, epidemiological and experimental evidence supports similar trans-generational effects^4,5^ although the mechanisms that underpin these are incompletely understood^6-9^. We generated a luciferase reporter knock-in mouse for the imprinted *Dlk1* locus, to visualise and track epigenetic fidelity across generations. We showed that exposure to high-fat diet (HFD) in pregnancy provokes sustained re-expression of the normally silent maternal *Dlk1* allele in offspring, coincident with increased DNA methylation at the *Dlk1 sDMR*. Interestingly, maternal *Dlk1* mis-expression was also evident in the next generation (F2), exclusively in animals derived from F1-exposed females. Oocytes from these females showed altered microRNA and gene expression, without any major changes in underlying DNA methylation, and correctly imprinted *Dlk1* expression resumed in subsequent generations (F3 onwards). Our results reveal how canonical and non-canonical imprinting mechanisms enable the foetal epigenome to adapt to *in utero* challenge to modulate the properties of two successive generations of offspring.

## Introduction and Results

Genomic Imprinting is an epigenetically regulated process that restricts mammalian gene expression in a parent-of-origin specific manner^10,11^. Mono-allelic gene expression is initiated by differential DNA methylation of parental germlines but is often reinforced post-fertilisation by the acquisition of additional epigenetic features that help sustain appropriate allelic expression (or silencing) within somatic tissues^12-14^. As a group, imprinted genes are critical for controlling embryonic growth and placental development^12,13,15^ and have key roles later in postnatal life, where they influence metabolism, neurogenesis and behaviour^16-18^. Expression of imprinted genes is tightly regulated, and subtle changes in expression often lead to profound changes in phenotype^16,19^. *Dlk1* is a prototypic paternally expressed, imprinted gene that is broadly expressed in the mid-gestation embryo but becomes increasingly restricted in the adult to subpopulations of cells in the adrenal and pituitary glands, skeletal muscle and brain^20-24^. Paternally restricted expression of *Dlk1* is associated with reciprocal expression of maternal *Gtl2 (Meg3), Rian (Meg8)*, anti-sense *Rtl1*, as well as clusters of intergenic microRNAs (*Mirg*) that collectively comprise one of the largest microRNA (miRs) clusters in the genome^25^. Molecular studies have shown that imprinting of the *Dlk1-Dio3* region is primarily regulated by a differentially methylated region (DMR), the *IG-DMR*, that shows selective methylation on the paternally inherited allele. Localised methylation across the *Dlk1 sDMR* and *Gtl2 sDMR* occur after fertilization and reinforce allelic marking to ensure expression of *Dlk1* and *Gtl2* from paternal and maternal alleles respectively^22,26,27^ (Figure 1A).

**Figure 1.**
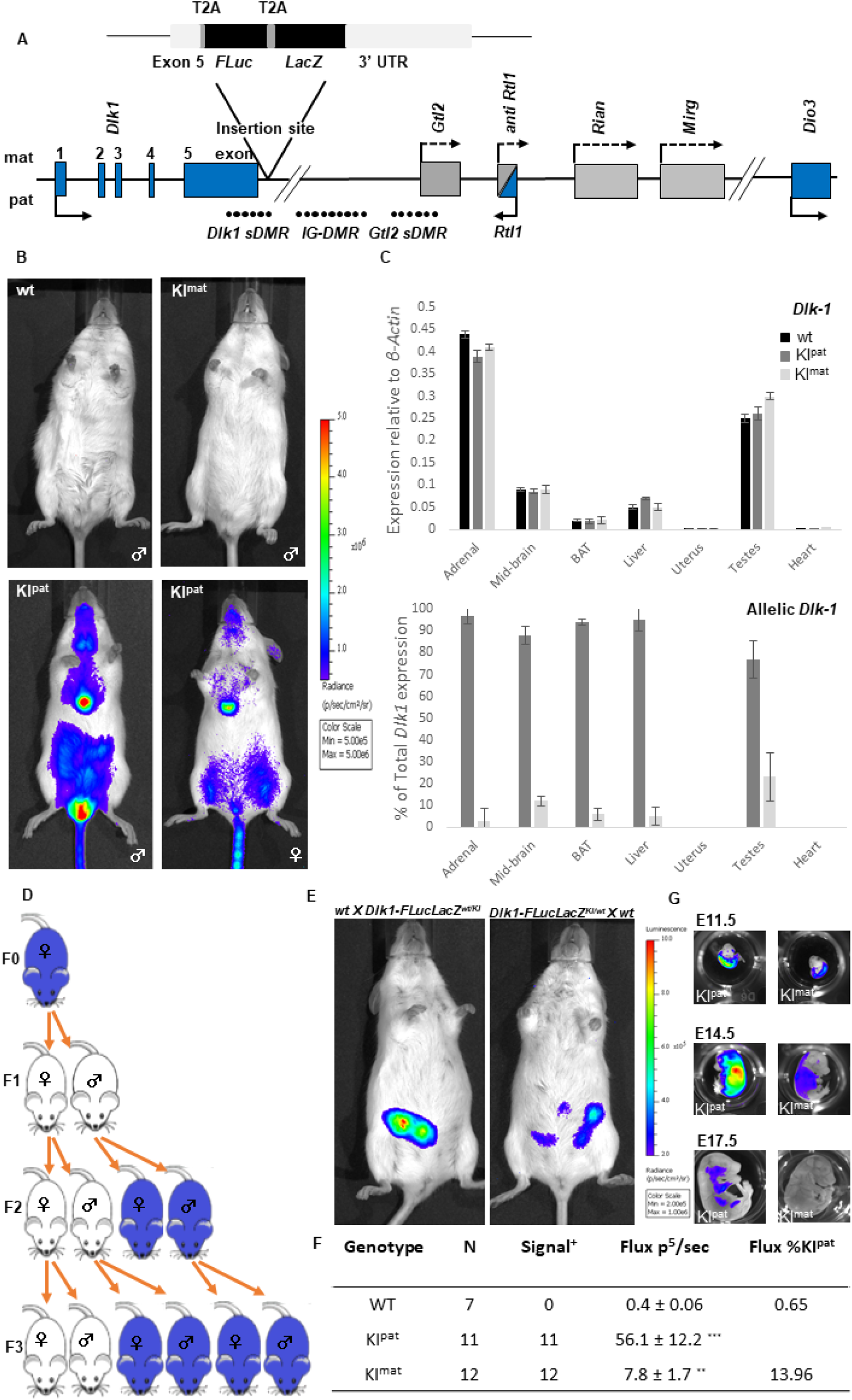
Generation and characterisation of reporter mice for imprinted *Dlk1* expression. A. Schematic of the mouse *Dlk1-Dio3* imprinted locus showing reporter insertion. Three differentially methylated regions (DMRs) that regulate imprinted expression of the cluster are indicated (closed circles represent methylated CpGs, *IG-DMR, Dlk1 sDMR* and *Gtl2 sDMR*) and the position of maternally expressed (light grey) and paternally expressed (blue) genes are shown. Arrows depict transcriptional direction, with solid lines representing protein coding genes and striped lines representing non-coding transcripts. In the *Dlk1-FLucLacZ* reporter line, *firefly Luciferase* (*FLuc*) and *β–galactosidase* (*LacZ*) were knocked into the endogenous *Dlk1* locus, with T2A sites downstream of exon 5. B. Bioluminescence (BL, blue) was detected in 8-week old male (lower panel, left) and female mice (lower panel, right) after paternal transmission of the reporter (KI^pat^). Minimal signal was detected in animals after maternal reporter transmission (KI^mat^, upper panel, right) or in wild type animals (wt, upper panel, left). Strong BL signal was evident in the thymus, central sternum and testes. C. *Dlk1* expression analysed by QRT-PCR (upper panel) was compared in different tissues from 8-week old male mice that inherited the reporter paternally (KI^pat^, dark grey), maternally (KI^mat^, light grey), or in non-transgenic controls (wt, black). Allelic *Dlk1* analysis (lower panel), using primers that distinguish the reporter from the wt allele showed a strong bias for paternal allele expression (dark grey) compared to maternal allele expression (light grey). (N: 4+4+4). D. Transmission of mono-allelic imprinted *Dlk1* reporter expression in four generations (F0, F1, F2, F3); upon paternal inheritance of the *Dlk1* reporter was expressed (blue), while maternal inheritance resulted in reporter silencing (white). Imprinting was predictably re-set across generations, through both germlines. (N: ≥2 independent litters per generation and reciprocal cross). E. BL signal (blue) detected in *Dlk1-FLucLacZ* pregnancies arising from KI^pat^ (left) and KI^mat^ (right) transmission, showed greater surface signal (E11.5) in KI^pat^ pregnancies. F. Quantification of BL signal (Flux) detected in E11.5 *Dlk1-FLucLacZ* embryos following dissection, demonstrating higher levels of signal from KI^pat^ than KI^mat^ embryos. BL signal in wt and KI^mat^ embryos, as a percentage of the average KI^pat^ signal is shown (N indicated, ^**^: P<0.01 ^***^: P<0.001.). G. BL imaging of embryos at different stages (E11.5, E14.5 and E17.5) showed progressive reduction in signals (blue) in both KI^pat^ (lower panel) and KI^mat^ (upper panel) through gestation; signal was readily detected in E11.5 and E14.5 KI^pat^ and KI^mat^ embryos, but at later stages (E17.5) was only seen after paternal transmission. Signal intensity scales are equalised between images.

Luciferase-based imaging offers a powerful non-invasive approach to visualise gene expression longitudinally in mammals^28^. By targeting luciferase into endogenous imprinted genes, such as *Cdkn1c*, we have been able to monitor allelic expression in living mice and demonstrate that exposure to chromatin-modifying drugs or dietary stress *in utero* can result in sustained loss of imprinting (LOI) in offspring^28^. To ask whether diet-induced deregulation of imprinting was heritable, we generated a mouse reporter in which *firefly luciferase* (*FLuc*) and *β-galactosidase* (*LacZ*) were knocked into the endogenous *Dlk1* locus (Figure 1A). To confirm luciferase activity accurately reports *Dlk1* expression in these mice, we performed a series of bioluminescence (BL) imaging and molecular analyses. The engineered *Dlk1* reporter allele retained faithful paternal expression^16,24^, with BL exclusively detected in *Dlk1-FLucLacZ* adult mice that inherited the reporter paternally (KI^pat^, Figure 1B) and signal localised to the brain, abdomen, testes, liver and a prominent signal corresponding to adrenal glands. No BL signal was seen in *Dlk1-FLucLacZ* mice inheriting the reporter maternally (KI^mat^) or in wild type control animals (wt). Tissue-specific and allelic *Dlk1* expression in the *Dlk1-FLucLacZ* reporter mice was verified by QRT-PCR. As shown in Figure 1C (upper panel), *Dlk1* transcript levels were similar in reporter and non-transgenic animals, consistent with minimal locus disruption. As anticipated, transgene-derived *Dlk1* expression was detected in adrenal, midbrain, liver, testes, and in brown adipose tissue (BAT), but not in uterine or heart tissue, and showed a strong bias in expression from the paternal allele (Figure 1C, lower panel). DNA methylation at the *Dlk1 sDMR, IG-DMR* and *Gtl2 sDMR* was comparable between reporter (*Dlk1-FLucLacZ* KI^pat^) and wild type controls (Figure S1A) and each of the four major isoforms of *Dlk1* were also detected at similar levels (Figure S1B). Taken together these results indicate that *Dlk1-FLucLacZ* accurately reports endogenous *Dlk1* expression and that reporter insertion had not substantially altered the methylation of regulatory DMRs. To verify that imprint erasure and re-setting occurs correctly in the *Dlk1-FLucLacZ* reporter line, we tracked BL activity in animals established from reciprocal crosses spanning four generations (N>6 per reciprocal cross and generation). Figure 1D illustrates these results, showing that epigenetic inheritance in the *Dlk1-FLucLacZ* colony followed the expected pattern for a paternally expressed imprinted gene. BL imaging studies revealed signal in the abdomen of pregnant mice carrying embryos with a paternally inherited *Dlk1*-reporter (Figure 1E, left) and dissection and *ex vivo* imaging confirmed expression in E11.5 embryos (Figures 1F and S2A). Transiently, from E11.5 to E14.5 a weaker signal was seen in embryos with a maternally inherited *Dlk1*-reporter (Figure 1E, right) but this was extinguished by E17.5 (Figure 1G). Temporal reinforcement of mono-allelic *Dlk1* expression during development is consistent with prior reports^29-31^ and was confirmed by whole-mount staining for *β-galactosidase* and optical projection tomography (Figure S2B).

Foetal exposure to maternal diet low in protein, or high in fat, can provoke long-lasting changes in gene expression, physiology and behaviour in offspring^4,32-34^. We showed previously that exposure to low protein diet *in utero* can provoke sustained LOI of the maternally expressed imprinted gene *Cdkn1c* in offspring^28^. To examine whether the paternally expressed *Dlk1* gene was sensitive to dietary challenge, we crossed *Dlk1-FLucLacZ* females with wild type males and exposed pregnant dams to either control diet (CD), low protein diet (LPD) or high fat diet (HFD) throughout pregnancy, as outlined in Figure 2A. Maternally transmitted *Dlk1-FLucLacZ* is predicted to be silent in offspring derived from these crosses. Consistent with this in all F1 animals that had been exposed to CD *in utero*, BL signal was near background. Imprinted expression was also maintained in all F1 animals that had been exposed to LPD *in utero*. In sharp contrast, exposure to HFD resulted in F1 animals (F1^mat-HFD^) that expressed maternally derived *Dlk1* (19/21, Figure 2B). This LOI was observed in mature male and female F1^mat-HFD^ offspring, long after gestational exposure and after being switched back to CD. LOI was also evident in HFD-exposed embryos at E17.5 (Figure 2C), at a time in gestation where *Dlk1* expression is shown to be exclusively derived from the paternal allele (Figure 1G). Expression of maternal *Dlk1* in HFD-exposed F1 animals showed a similar tissue distribution as paternally-derived *Dlk1* (in brain, liver, thyroid, BAT and testes) with the exception of ectopic expression in the uterus (Figure 2D). LOI was associated with a selective increase in DNA methylation at *Dlk1 sDMR* and *Gtl2 sDMR* (but not *IG-DMR*) in the affected tissues (liver and in BAT, exemplified in Figure 2E) and with an increase in gene expression across the entire gene cluster (Figure 2F, Figure S3A).

**Figure 2.**
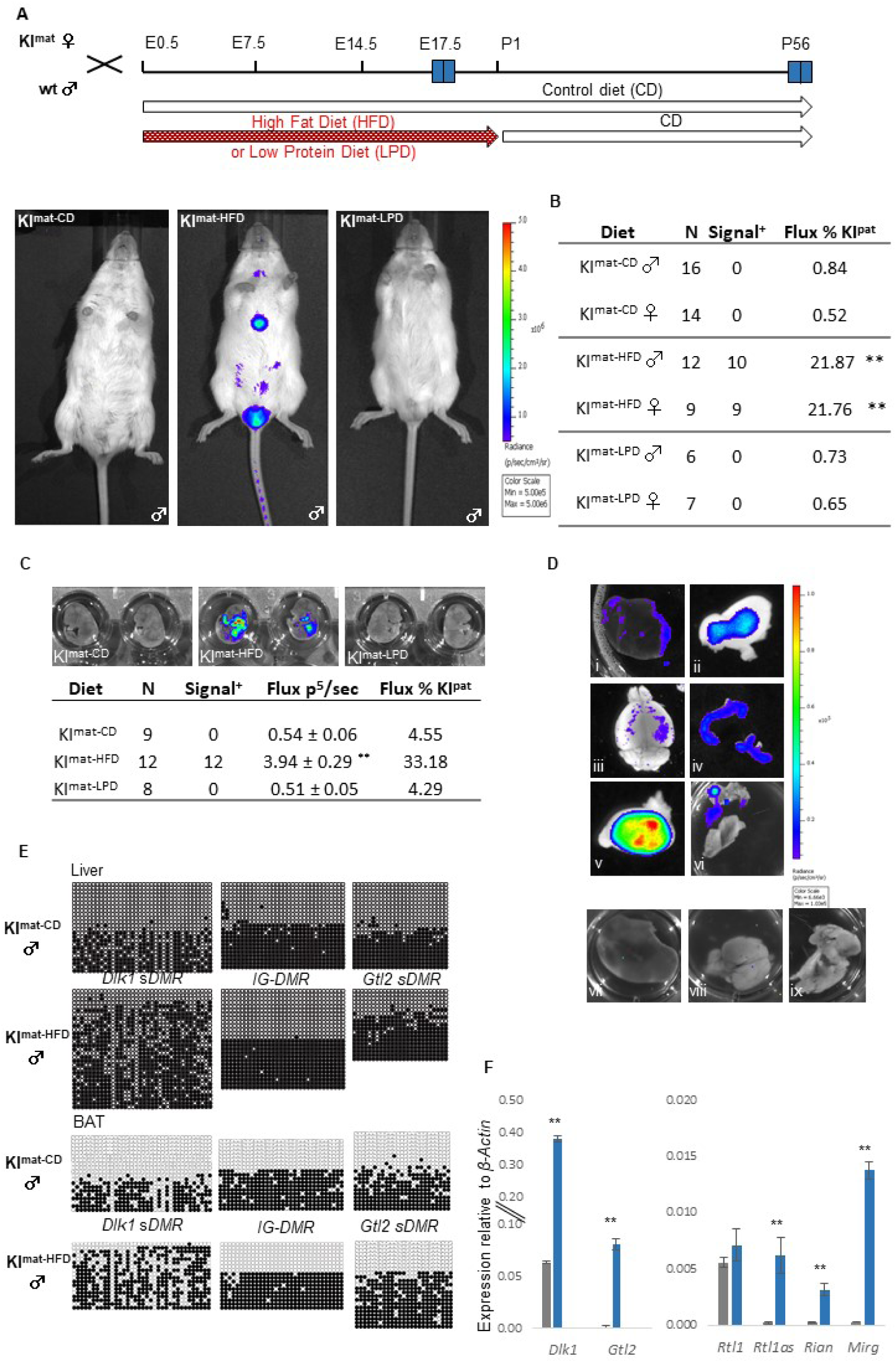
Exposure to high fat diet *in utero* results in loss of *Dlk1* imprinting in offspring. A. Schematic of experimental breeding, dietary regime and bioluminescent imaging. Offspring inheriting *Dlk1-FLucLacZ* maternally (KI^mat^) were generated by mating wt males with heterozygous *Dlk1-FLucLacZ* females; upon detection of a vaginal plug pregnant females were maintained on a control (CD) diet, or switched to a calorie-matched low protein diet (LPD) or a high fat diet (HFD) for the duration of the pregnancy. At birth, all animals were maintained on CD and BL imaging was performed at the times indicated (E17.5 and post-natal day 56). Increased BL signal (blue) was evident in 8-week old mice that had been exposed to gestational HFD (F1^mat-HFD^, middle image), as compared to either CD or LPD-exposed animals (KI^mat-CD^ left and right, respectively). B. Quantification of abdominal bioluminescence showed a significant increase in signal in F1^mat-HFD^ offspring as compared to KI^mat-CD^ or F1^mat-LPD^. BL signal in F1^mat-HFD^ animals was less than that in dietary control animals that inherit the reporter paternally (KI^pat-CD^), suggesting a partial release of silencing. (P<0.01 **). C. BL signal (blue) in E17.5 embryos from F1^mat-HFD^, F1^mat-LPD^ and KI^mat-CD^ are compared (upper panel) and quantified (lower panel showing Flux levels relative to KI^pat-CD^ controls). BL signals were significantly higher in F1^mat-HFD^ than F1^mat-LPD^ and KI^mat-CD^ embryos. (P<0.01 **). D. BL signal (blue) detected *ex vivo* in organs of 8-week old F1^mat-HFD^ animals (upper panels: i-liver, ii-white adipose, iii-brain, iv-uterus, v-testes, vi-brown adipose tissue). Control tissues from 8-week old KI^mat-CD^ animals (lower panel: vii-liver, viii-brain, ix-brown adipose) are shown for comparison. E. DNA methylation analysis at the *Dlk1 sDMR, IG-DMR* and *Gtl2 sDMR* in liver (upper) and brown adipose tissue (BAT) (lower) from 8-week old KI^mat-CD^ and F1^mat-HFD^ animals. Hyper-methylation was detected at the *Dlk1 sDMR* (CD 31.1 ± 5.8 % vs. HFD 67.3 ± 9.5 %, P<0.01 **) and the *Gtl2 sDMR* (CD 44.7 ± 5.3 % vs. HFD 64.2 ± 7.4 %, P<0.05 *) in F1^mat-HFD^ liver, but the *IG-DMR* (CD 51.7 ± 2.2 % vs. HFD 49.7 ± 4.8 %, NS) was unchanged. Similar results were seen for BAT. Closed circles indicate methylated CpG and open circles un-methylated CpG. Each row represents an individual clone. (N: 4+4 with 2 KI^mat^ and 2 wt animals per group). F. Comparison of gene expression at the *Dlk1-Dio3* cluster, analysed by QRT-PCR in liver of 8-week old F1^mat-HFD^ (blue) and KI^mat-CD^ (dark grey). (N: 4+4, P<0.01 **). Double lines represents a scale break.

To investigate whether HFD-induced alteration of *Dlk1* was transmitted to subsequent generations, we examined F2 mice derived from crosses between F1^mat-HFD^ females and wild type males (F2^mat-HFD^, Figure 3A). In this setting maternal inheritance is predicted to ensure *Dlk1* silencing, however BL signal was detected in most F2^mat-HFD^ offspring, as illustrated for a litter of eleven animals (Figure 3B) in which seven were transgenic (KI^mat^). BL signal distribution suggested that *Dlk1* mis-expression was variable among F2^mat-HFD^ animals but lower than signal seen in KI^pat^ controls (Figure 3C), consistent with partial LOI. Molecular analyses confirmed heterogeneity (Figure S4) and ectopic expression (Figures 3D) of maternally-derived *Dlk1* among F2^mat-HFD^ male and female offspring. Despite partial loss of *Dlk1* imprinting, overall levels of *Dlk1* expression in F2^mat-HFD^ were generally lower in adrenal, midbrain, BAT and testes (Figure 3D, red) than in controls (F2^mat-CD^, black), with the notable exception of uterine tissue. In somatic tissue, increased DNA methylation at the *IG-DMR* and *Dlk1 sDMR* and decreased methylation at the *Gtl2 sDMR* was evident in F2^mat-HFD^ samples (Figure 3E), consistent with altered expression of maternal *Dlk1*. In contrast, F2 offspring generated from reciprocal crosses (F1^mat-HFD^ males x wild-type females) showed *Dlk1* reporter expression consistent with normal paternal inheritance and imprinting (Figure 3C, litters 7 and 8).

**Figure 3.**
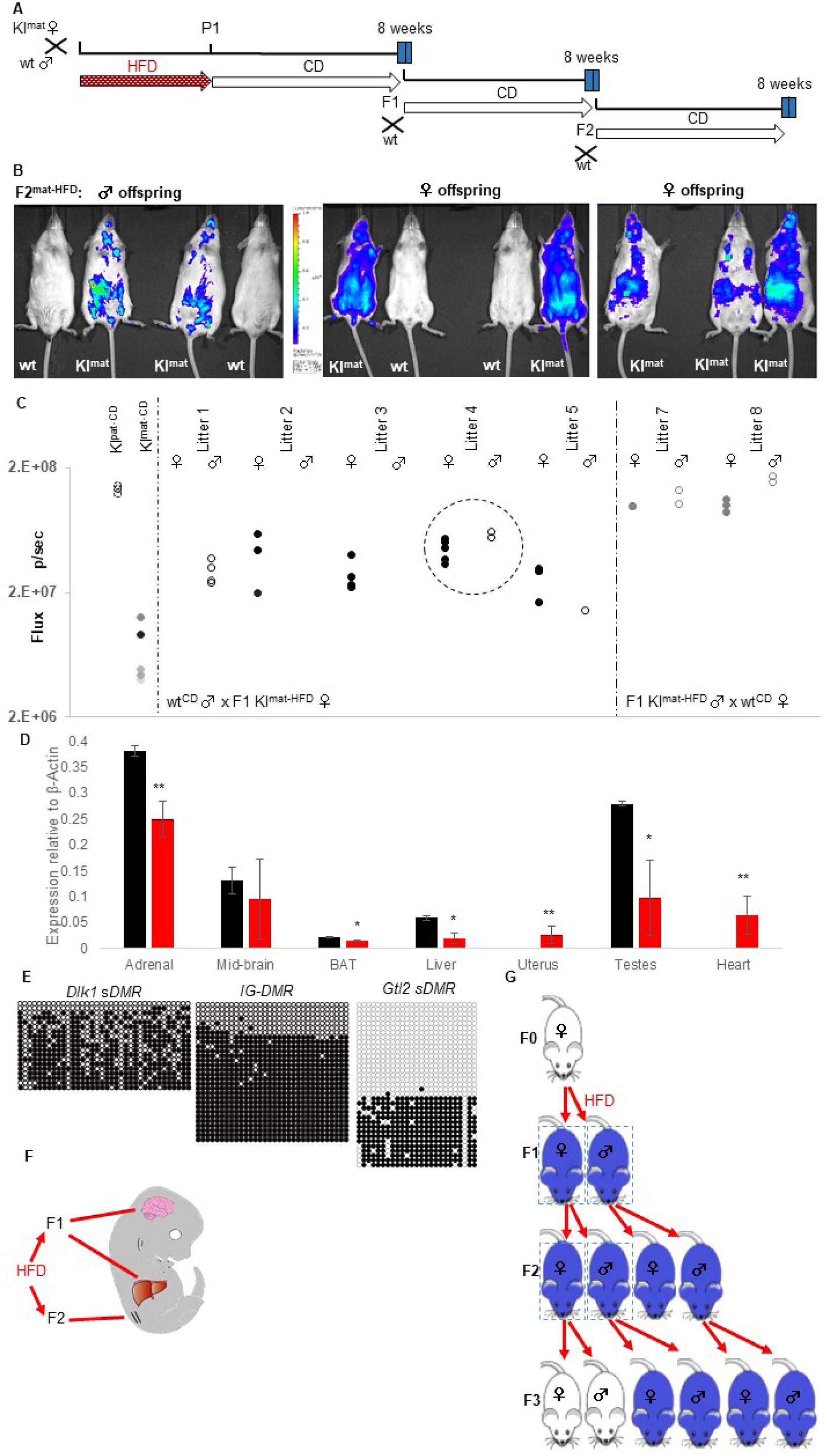
Exposure-induced changes to *Dlk1* imprinting are transmitted to F2 offspring. A. Schematic of generational studies of the impact of *in utero* dietary exposure. Animals that had been exposed to gestational HFD (*Dlk1-FLucLacZ* F1^mat-HFD^) were imaged and at 8 weeks of age mated with a wild type (CD-fed) mate. Dams and offspring were maintained on CD for the duration of their pregnancy, lactation and weaning and offspring (F2 and F3) were examined at the times indicated. B. BL signal (blue) was detected in F2 offspring (F2^mat-HFD^) derived from F1 females that had been exposed to gestational HFD. In the affected F2^mat-HFD^ animals, BL signal distribution and intensity was variable and ectopic. C. Quantification of abdominal BL signal (Flux p/sec in area of interest) in individual 8-week old F2^mat-HFD^ animals, generated from 6 individual F1^mat-HFD^ females and wt^CD^ males (litters 1-5, no litters generated by female 6) or 2 individual F1^mat-HFD^ males and wt^CD^ females (litter 7,8). BL signal intensity in KI^mat-CD^ and KI^pat-CD^ animals provide controls for comparison. Highlighted litter (4) is represented in (B). D. QRT-PCR analysis showing *Dlk1* expression in adrenal, mid-brain, BAT, liver, and testes from F2^mat-HFD^ animals (red) was lower than in KI^mat-CD^ samples (black bars). In F2^mat-HFD^ animals *Dlk1* expression was detected in the uterus and heart, organs which do not normally express *Dlk1* (N: 4+4) (P <0.05 *, P <0.01 **). E. DNA methylation analysis at the *Dlk1 sDMR, IG-DMR* and *Gtl2 sDMR* in liver of 8-week old F2^mat-HFD^ mice showed hyper-methylation at the *Dlk1 sDMR* and *IG-DMR*. Closed circles indicate methylated CpG and open circles un-methylated CpG. Each row represents an individual clone. Images are representative. (N: 3). F. Hypothesis that multi-generational impact (on F1 and F2) can arise from a single gestational exposure. G. Summary of alterations in *Dlk1* expression induced by gestational exposure to HFD in pregnant F0 females. Normally *Dlk1* is silent (white) when transmitted maternally. Gestational exposure to HFD in F0 females provokes loss of imprinting in F1 offspring (blue, box). F1 females transmit altered *Dlk1* expression to their offspring (F2 blue, box). F1 males and F2 females transmit *Dlk1* appropriately so that in F3 mice, *Dlk1* expression is parent-of-origin specific.

One explanation for the LOI in second-generation offspring from HFD-exposed females is that *in utero* exposure affects the developing epigenome of oocytes contained within developing female embryos, as well as somatic tissue (Figure 3F). In this scenario a trans-generational mechanism of epigenetic inheritance is not strictly required^7^ and imprinting would be predicted to resume in subsequent generations (Figure 3G). Consistent with this, BL signal was undetectable in all F3 transgenic animals derived from LOI-affected F2^mat-HFD^ dams (Figure S5).

To better understand the basis of ectopic *Dlk1* reporter expression in F2^mat-HFD^ offspring, we asked whether DNA methylation was perturbed in the gametes of F1^mat-HFD^ mice. As predicted, F1^mat-HFD^ sperm showed DNA methylation exclusively at the *IG-DMR*, with hypo-methylation at *Dlk1 sDMR* and *Gtl2 sDMR* (Figure S6). F1^mat-HFD^ oocytes were collected individually and processed for parallel genome-wide single-cell bisulphite sequencing and single-cell RNA-seq (scM&Tseq)^35,36^. In both groups the anticipated bimodal pattern of DNA hypo- and hyper-methylated domains in oocytes^36,37^ was retained, with a broadly similar profile assessed over 100-CpG windows (r=0.984) (Figures S7A, S7B). While differences in methylation were detected in 439 differentially methylated 100-CpG tiles (representing approximately 0.2% of genomic tiles) these were dispersed across the genome (Figure S7C) and there was no obvious separation of oocytes from different groups from principal component analysis (Figure S7D). Among the imprinted gametic DMRs, high levels of methylation of maternal gDMRs and low level of methylation of paternal gDMRs were well preserved in F1^mat-HFD^ oocytes (Figure 4A) and we did not detect significantly increased variation or anomalous gDMR methylation in HFD-exposed oocytes as compared with CD (Figure S7E). At the *Dlk1-Dio3* imprinted domain itself, the *IG-DMR* domain remained similarly hypo-methylated in CD and HFD groups, while the *Dlk1 sDMR* and *Gtl2 sDMR* showed modestly increased methylation in HFD oocytes (Figure 4B). The lack of statistically significant changes in DNA methylation levels at the *Dlk1-Dio3* imprinted domain make it unlikely that this is responsible for the highly penetrant maternal transmission of ectopic *Dlk1* expression seen after *in utero* HFD exposure.

**Figure 4.**
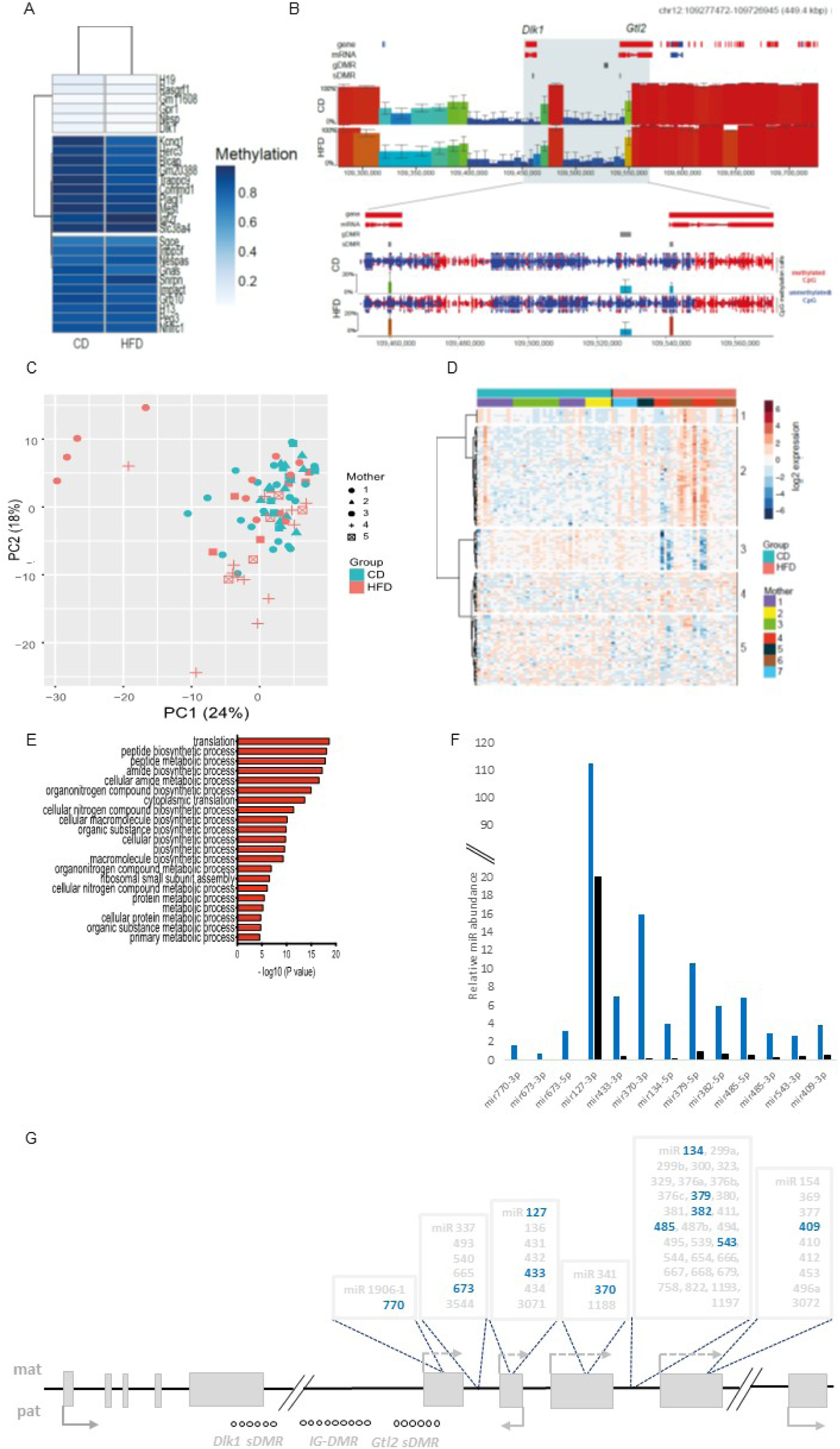
Germline DMRs in single MII oocytes from F1 females are unaffected by dietary exposure but show an altered transcriptional programme. A. Heatmap representing mean DNA methylation for each gametic (g)DMR in the CD and HFD groups (merged from 40 and 37 oocyte scBS-seq datasets, respectively). B. SeqMonk screenshot showing mean DNA methylation in CD and HFD oocytes over non-overlapping 100 CpG windows. Zoomed in region showing the CpG methylation calls (methylated red; un-methylated, blue) of the *Dlk1-Dio3* imprinted cluster. C. Principal component analysis of scRNA-seq datasets of individual oocytes from F1 females exposed *in utero* to HFD and CD controls. D. Heatmap revealing 5 unsupervised clusters of the 198 most variable genes. Top bars identify the F1 donor and diet groups. E. Major terms highlighted in the gene ontology analysis of up-regulated genes from clusters 1, 2 and 4 (-log10 of adjusted P value). F. Comparison of *Dlk1-Dio3* microRNAs (miRs) detected in HFD-exposed F1 oocytes (blue) versus CD-exposed F1 oocytes (black) (N:4+4). All represented miRs were found to be significantly more represented in F1^mat-HFD^ oocytes (P<0.01). Double line represents a scale break. G. Schematic showing miR species upregulated in HFD-exposed F1 oocytes (blue) span multiple domains within the *Dlk1-Dio3* miR cluster.

Genome-wide transcription was also examined in these F1^mat-HFD^ oocytes. Although PCA indicated no clear separation of oocytes according to CD and HFD groupings, increased heterogeneity was evident in the HFD group (Figure 4C) and a subset of 198 genes showed highly variable expression (>0.528 fold from the mean standard deviation (Figure S8A). These genes also showed the largest differences in expression between groups (Figure S8B). Unsupervised hierarchical clustering segregated the variable genes into five clusters (Figure 4D), the largest of which (87 genes), that were more likely to be upregulated in HFD oocytes, were associated with regulation of translation and modulation of biosynthetic and metabolic processes (Figure 4E). Therefore, in marked contrast with the very limited differences seen in DNA methylation, our results indicate altered transcriptional quality of oocytes from *in utero* HFD-exposed F1 females. Importantly, a marked increase in the expression of multiple miR species in F1 oocytes that had been exposed to HFD as compared to CD was evident (Figure 4F). Up-regulation of these miRs was sustained for long periods after gestational exposure and affected clusters spanning the entire *Gtl2-antiRtl1-Rian-Mirg* domain (Figure 4G). These data show that microRNA expression in the developing oocytes is irrevocably altered in response to *in utero* exposure to HFD and implicate this in the subsequent mis-regulation of *Dlk1* in F2 progeny. Sustained disruption in expression across the *Dlk1-Dio3* domain was also evident in somatic tissue from F1^mat-HFD^ (Figure S3), F2^mat-HFD^ and F3^mat-HFD^ mice (Figure S5B) long after HFD exposure.

## Discussion

Our data show that dietary challenge during pregnancy induces phenotypic heterogeneity that impacts two generations of offspring. In F1 animals, high fat maternal diet induced a sustained loss of *Dlk1* imprinting in somatic tissue that was associated with increased methylation of the *Dlk1 sDMR*. In F2 animals born exclusively to F1-exposed females, expression of *Dlk1* was ectopic, implicating epigenetic alteration of the developing F1 oocyte epigenome. Female gametogenesis initiates in growing oocytes in the ovary post-natally^38-40^, so a direct effect of exposure on *de novo* DNA methylation is very unlikely. However, as exposure is contemporary with DNA methylation erasure in primordial germ cells, we asked whether de-methylation was impaired in F1-exposed oocytes, but found no evidence of either residual methylation of paternal gDMRs or altered DNA methylation across the *Dlk1-Dio3* domain. Instead, HFD-exposed F1 oocytes showed altered gene expression detected using genome-wide scRNA-seq, as well as pronounced changes in the expression of many *Dlk1/Gtl2*-associated genes and miRs. This result was surprising given that oocytes were sampled from F1 adults after ovulation and several weeks after HFD exposure, and implicates an alternative (non-canonical imprinting) route of intergenerational epigenetic transmission where chromatin and miR expression, rather DNA methylation, predispose to *Dlk1* and *Gtl2* deregulation. As histone H3K27me3 is pervasive in the oocyte genome and occupies DNA hypo-methylated domains^41,42^, and has been shown to be responsible for a parallel imprinting mechanism^43,44^, it is tempting to speculate that this modification may play a role. In oocytes and somatic tissue from F1-exposed animals, we found dramatic increases in *Gtl2, Rtl1as, Rian* and *Mirg* expression, in addition to a plethora of miRs that normally are expressed only at low levels and exclusively from the maternal allele^22,25,45^. Importantly, we show that this high level of transcription across the *Dlk1-Dio3* domain persisted in F1 animals even after being returned to a normal diet. These results suggest an alternative explanation for *Dlk1* deregulation seen in the next generation. As expression of miRs across the *Gtl2* domain may be self-sustaining^46-49^ and disruption of maternally expressed miRs can alter paternal *Dlk1* expression^49^, constitutive activity of this domain in oocytes is likely to result in an imbalance of *Dlk1* expression in F2 offspring^21,50-52^. Together these results raise an intriguing possibility that genomic imprinting mechanisms that harness multiple types of epigenetic control enable the phenotype of successive generations of offspring to be modified, in response to environmental challenges precociously sampled ahead of birth.

---

**Supplementary Figure 1.**
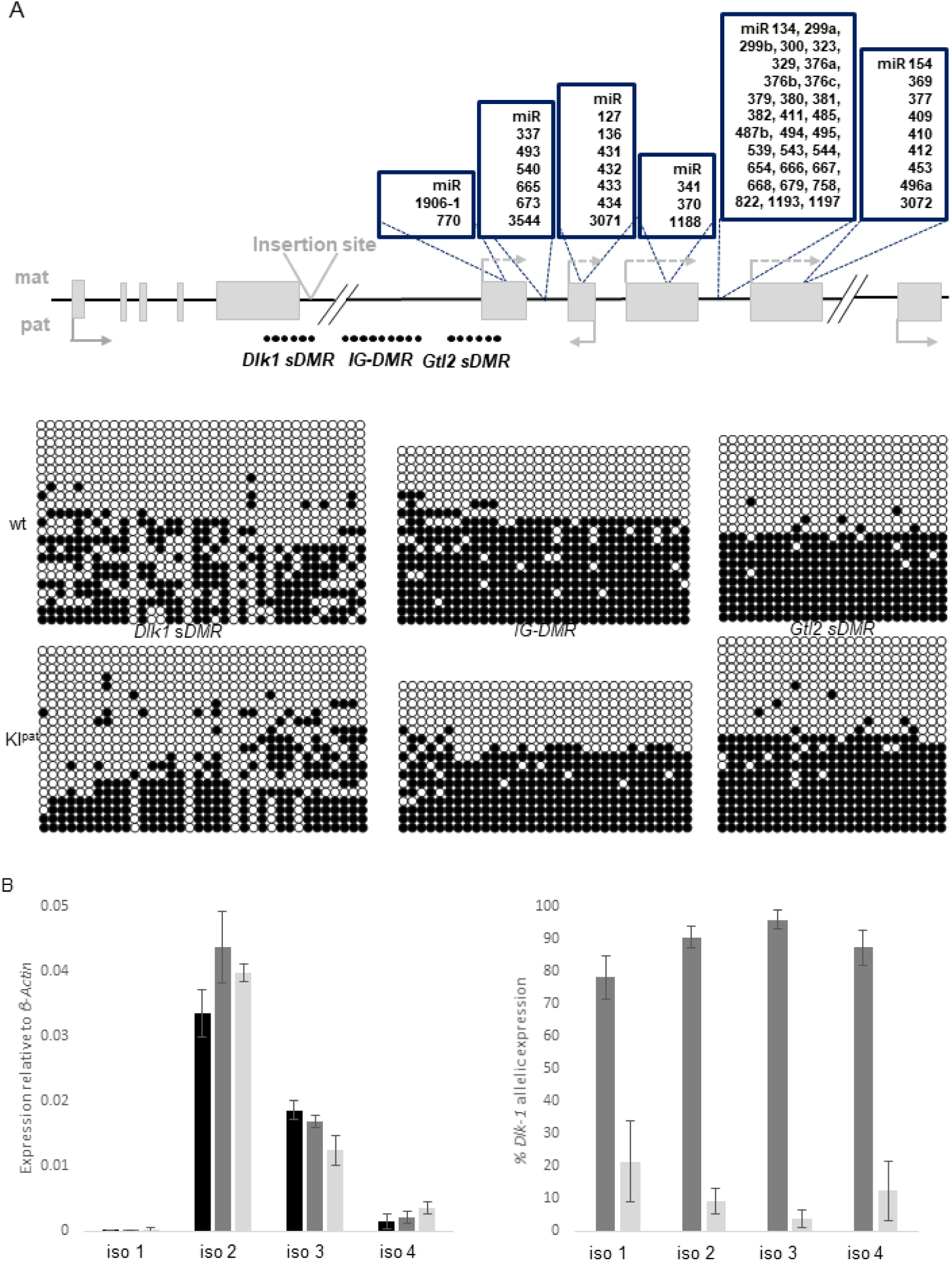
Knock-in reporter does not disturb *Dlk1* expression or imprinting. A. Expanded schematic for *Dlk1-Dio3* mouse imprinted region with scheme for knock-in strategy. The three differentially methylated regions (DMRs) that regulate imprinted expression of the cluster (*IG-DMR, Dlk1 sDMR* and *Gtl2 sDMR*) are depicted by closed circles. Methylation patterns of all three DMRs were found to be unchanged between wt and KI^pat^ 8-week old adult liver (closed circles-methylated CpG, open circle-un-methylated CpG. Each row represents an individual clone. N: 2+2). Arrows depict transcriptional direction, with solid lines representing protein coding genes, and striped lines representing non-coding transcripts. Numerous miRs are located within this cluster, highlighted above each location in ascending order. B. QRT-PCR of *Dlk1* isoforms (left graph) in 8-week old male liver reveals differential expression of individual isoforms, with the majority of *Dlk1* total expression contributed by isoforms 2 and 3. No differences were observed between wt (black bar), KI^pat^ (dark grey) and KI^mat^ (light grey) groups. Allelic expression averaged from both KI^mat^ and KI^pat^ samples, revealed a strong paternal bias in each isoform, with very low contribution from the maternally derived allele. (N: 4+4+4).

**Supplementary Figure 2.**
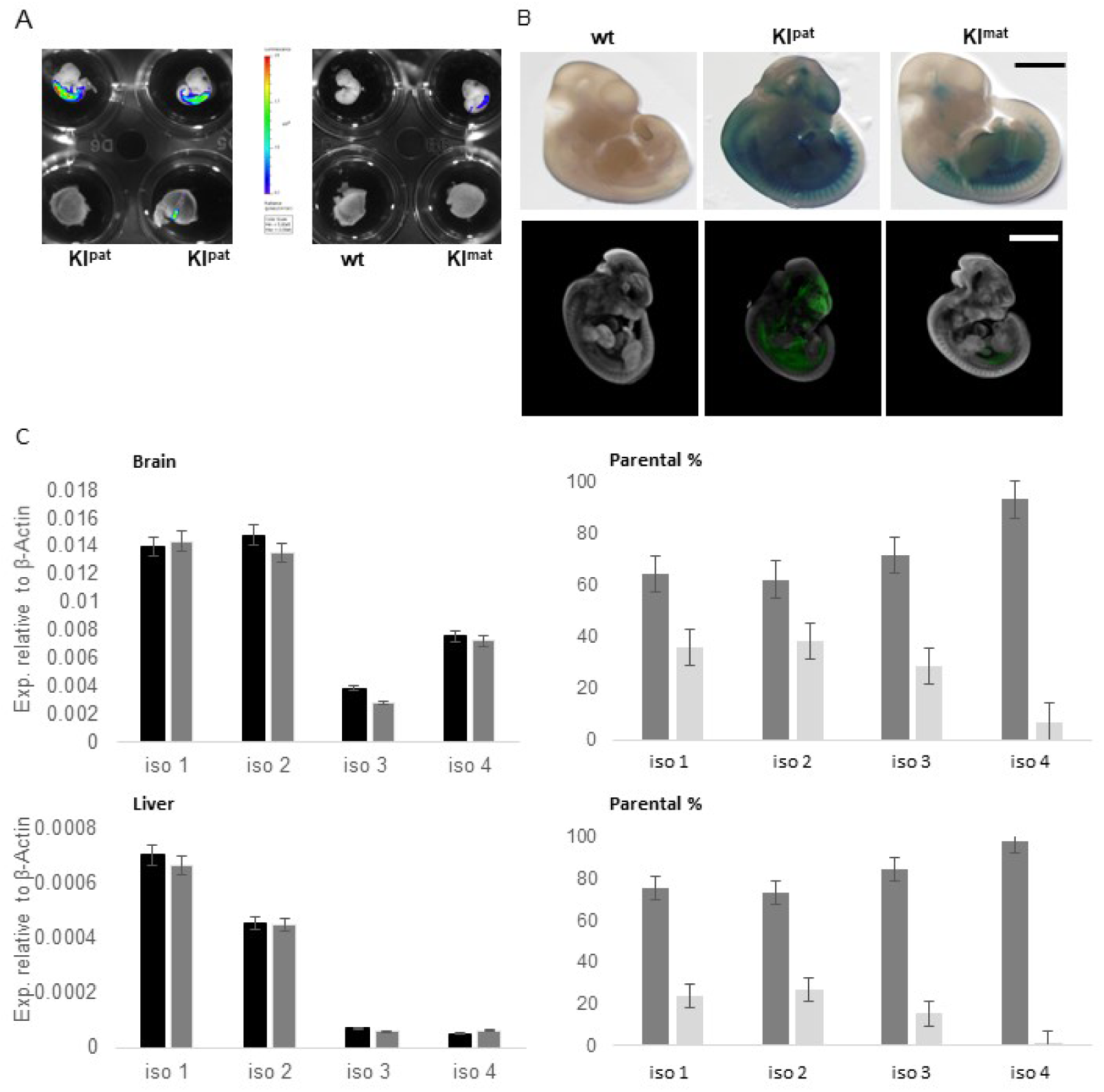
Overlap in expression of *Dlk1* maternal and paternal alleles in E11.5 embryos. A. BL imaging of *Dlk1-FLucLacZ* embryos at E11.5, where luciferase activity (blue) was seen in head and abdomen of both KI^pat^ and KI^mat^ embryos. B. LacZ staining of *Dlk1-FLucLacZ* whole embryos at E11.5 showing labelling of cartilage, brain and abdomen upon paternal inheritance of the reporter, with limited staining detectable upon maternal inheritance in the abdomen and no staining detected in wt embryos. Scale bar: 2 mm. Optical Projection Tomography (OPT) of LacZ stained E11.5 *Dlk1-FLucLacZ* embryos. Absorbance (green) was measured in the liver, cartilage, gonadal ridges and a subset of forebrain regions in KI^pat^ embryos. Absorbance was weaker in KI^mat^ embryos, with very low signal level detected in wt embryos. Scale bar: 2 mm. Images shown are representative. C. QRT-PCR analysis of *Dlk1* expression in brain (upper panel) and liver (lower panel) of E11.5 embryos. Isoform (left) contribution was unchanged in KI^pat^ (dark grey) compared with wt controls (black) in brain and liver. Similar levels of each of the four *Dlk1* isoforms was seen in brain, while expression in the liver was primarily restricted to isoforms 1 and 2. A significant contribution from the maternal *Dlk1* allele (light grey) was detected in brain and liver (right hand graphs), when compared to the paternal allele (dark grey), representing isoforms 1 and 2 with a much lower contribution of isoforms 3 and 4. (N: 4+4).

**Supplementary Figure 3.**
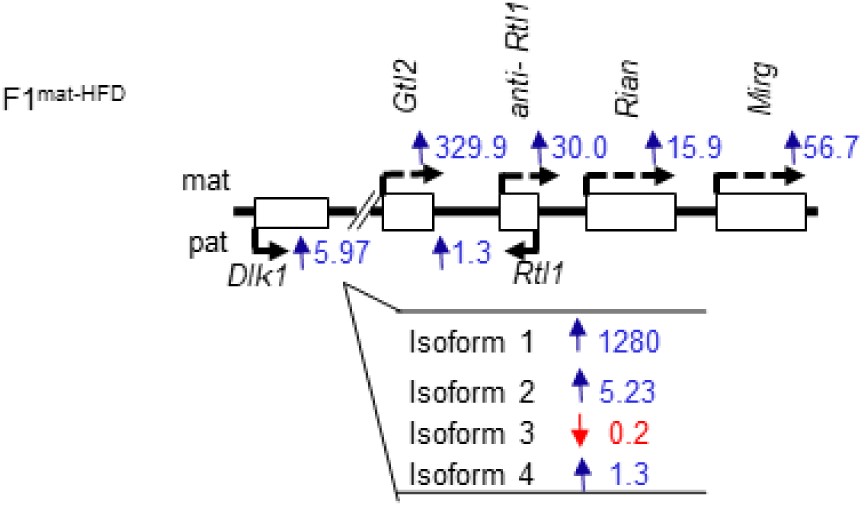
Expression of the *Dlk1-Dio3* cluster is altered in F1 and F2 animals after gestational HFD exposure. Changes in the expression of the *Dlk1-Dio3* cluster in F1^mat-HFD^ as determined by QRT-PCR analysis, where blue and red arrows indicate fold-increase and fold-decrease in transcript levels respectively in the liver of 8 week old mice, as compared to F1^mat-CD^ controls. Expression of all genes increased in F1^mat-HFD^ liver, with large increases in maternally-derived transcripts (such as *Gtl2, anti-Rtl1, Rian, Mirg*) that are normally expressed only minimally. Isoform-specific analysis of *Dlk1* revealed that elevated *Dlk1* expression was attributable to increased isoform 1 and 2 expression in F1^mat-HFD^. (N: 4+4).

**Supplementary Figure 4.**
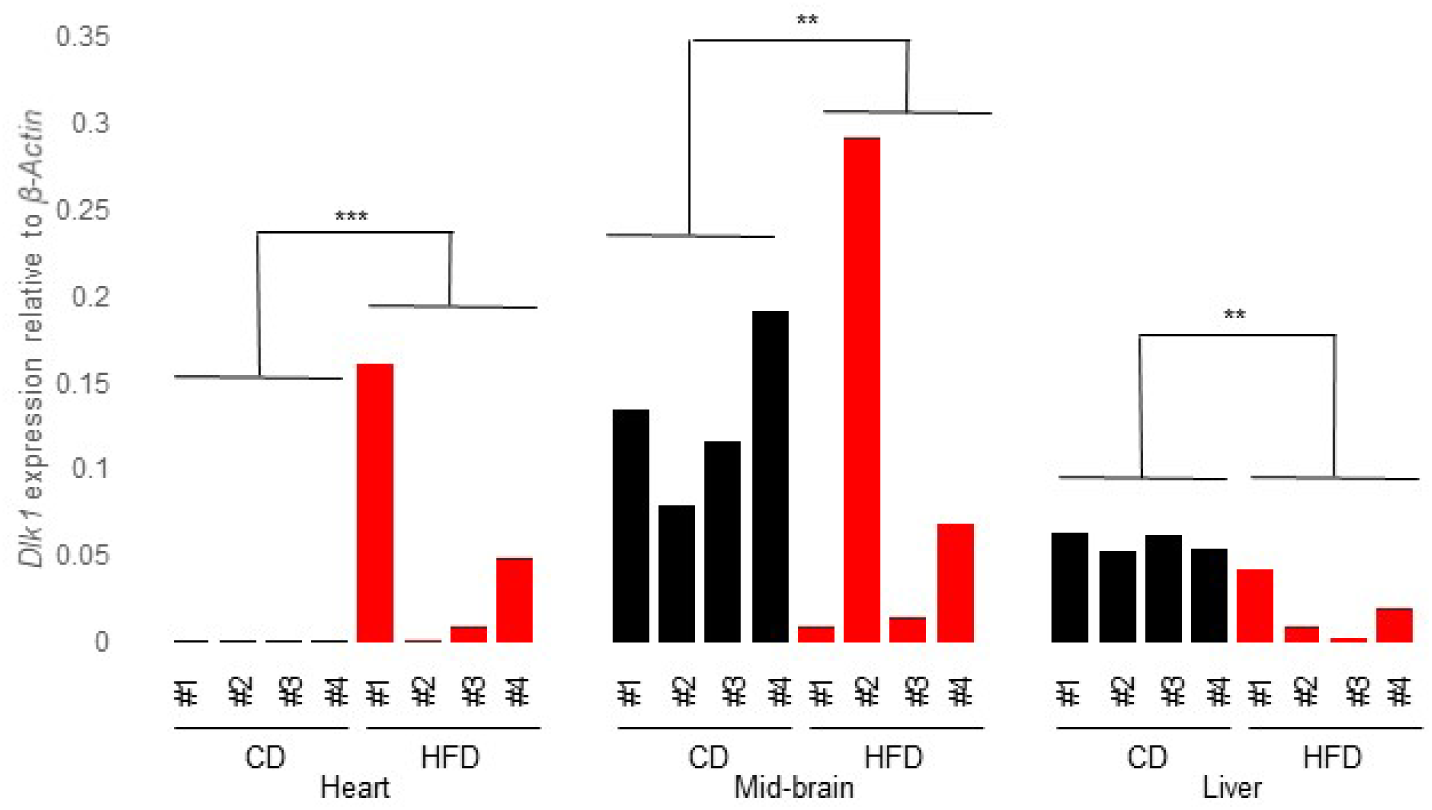
Ectopic and heterogenous expression of *Dlk1* in F2^mat-HFD^ animals. *Dlk1* expression in organs from individual control (black) and F2^mat-HFD^ (red) animals, as determined by QRT-PCR analysis. Highly variable expression levels were observed within tissue between samples of F2^mat-HFD^ animals, which was not found in control animals. (P< 0.01 **, P< 0.001 ***) (Variance Heart: CD 1.013×10^−10^, HFD 4.12×10^−2^, Mid-brain: CD 1.66×10^−3^, HFD 1.33×10^−2^, Liver: CD 2.61×10^−5^ HFD 2.17×10^−4^) (N: 4+4).

**Supplementary Figure 5.**
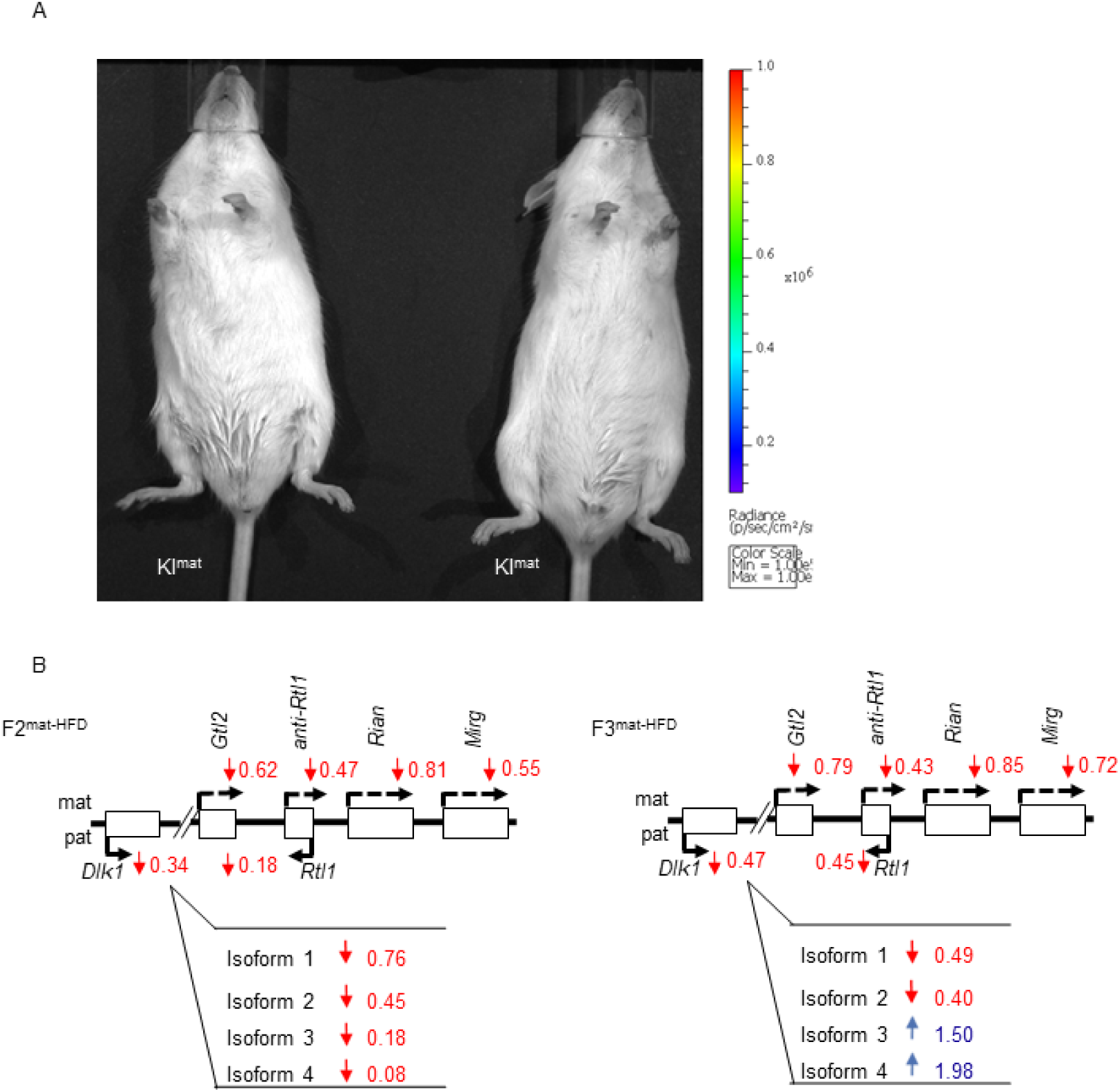
Imprinted *Dlk1* expression is largely restored in the F3 generation. A. BL imaging of *Dlk1-FLucLacZ* F3^mat-HFD^ 8-week old male mice, where minimal luciferase activity was detected (comparable with KI^mat-CD^ animals). B. Gene expression analysis of the *Dlk1-Dio3* cluster in F2^mat-HFD^ (left) and F3^mat-HFD^ (right) liver showing a generalised reduction in transcript levels (red arrows). (N: 4 per group.)

**Supplementary Figure 6.**
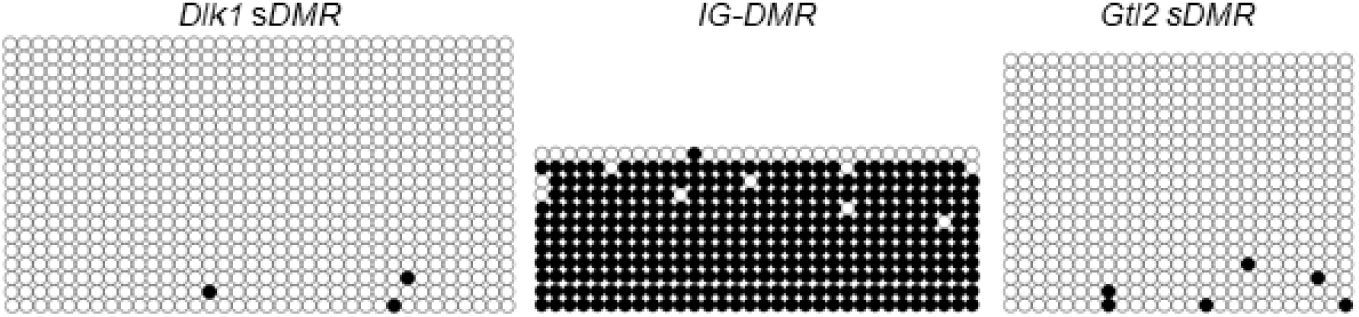
Methylation of *Dlk1-Dio3* cluster DMRs is unchanged in F1mat-HFD sperm. Bisulphite analysis showing DNA methylation at *Dlk1 sDMR* (left), *IG-DMR* (middle) and *Gtl2 sDMR* (right) in F1^mat-HFD^ 8-week old sperm. The *IG-DMR* was found to be hyper-methylated, while both the *Dlk1 sDMR* and *Gtl2 sDMR* were found to be largely un-methylated. (Closed circles-methylated CpG, open circle-un-methylated CpG. Each row represents an individual clone. N: 2, Images are representative).

**Supplementary Figure 7.**
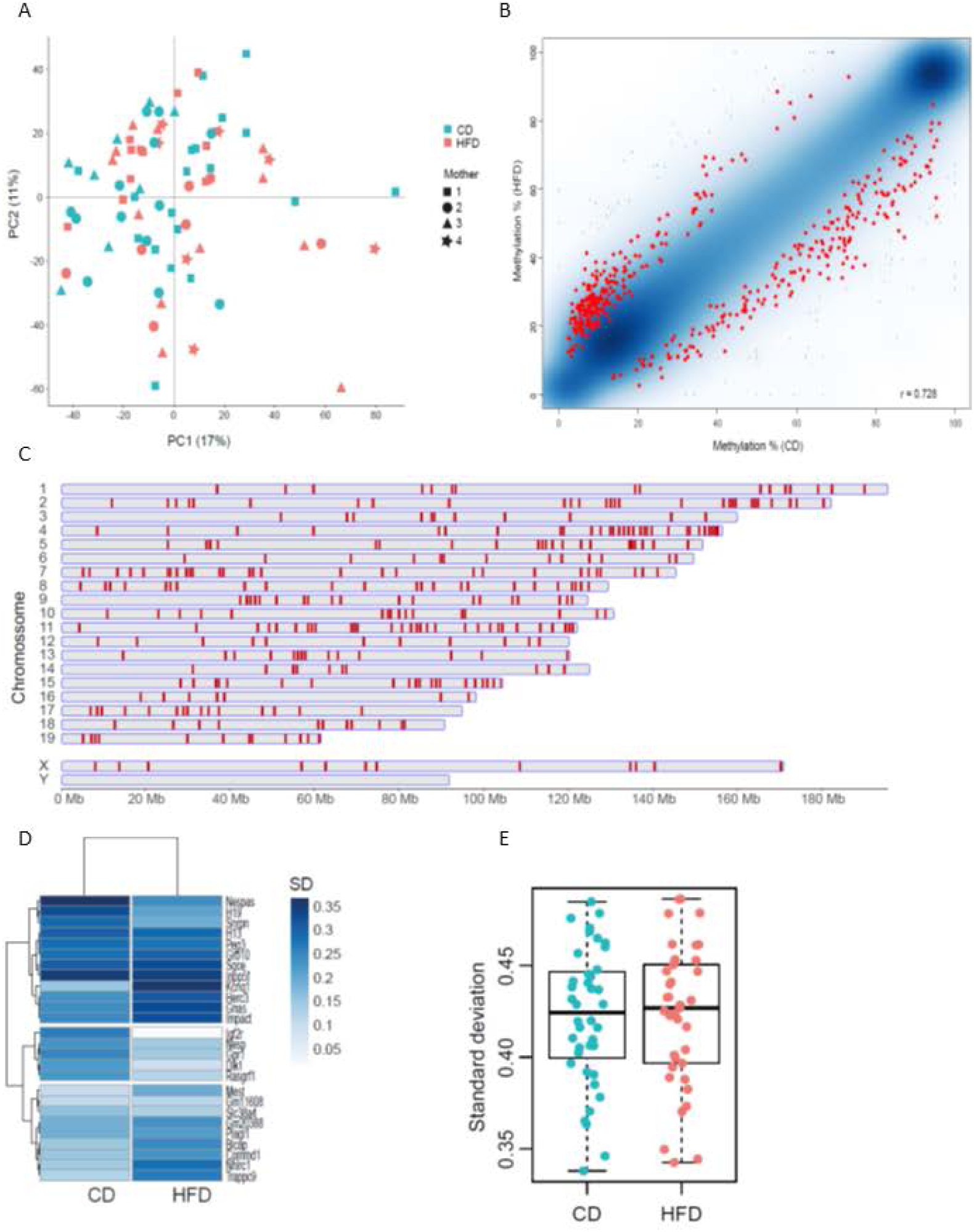
scDNA-methylation analysis of F1^mat-HFD^ oocytes. A. PCA plot of the 40 and 37 scBS-seq datasets of oocytes from CD and HFD F1 females, respectively. The plot is based on 20kb running windows with 2kb spacing informative in all scBS-seq datasets. B. Scatterplot of grouped data demarking the 459 100-CpG tiles (red) called differentially methylated between HFD and CD F1 oocytes with an absolute methylation difference of ≥10%. The 459 DMRs represent DMRs identified as significant at a FDR of <0.05, with at least 10% difference in methylation between groups, in 70% of 100 permutations of 36 cells in pools of 9 cells per group. C. Chromosome view showing distribution of DMRs D. Heatmap representing the variation (SD) in methylation of each gDMR across all the oocytes in the CD and HFD groups. E. Box-whisker plot representing the variation (SD) of all the informative gDMRs in each individual oocyte in the CD and HFD groups; each point represents a single oocyte.

**Supplementary Figure 8.**
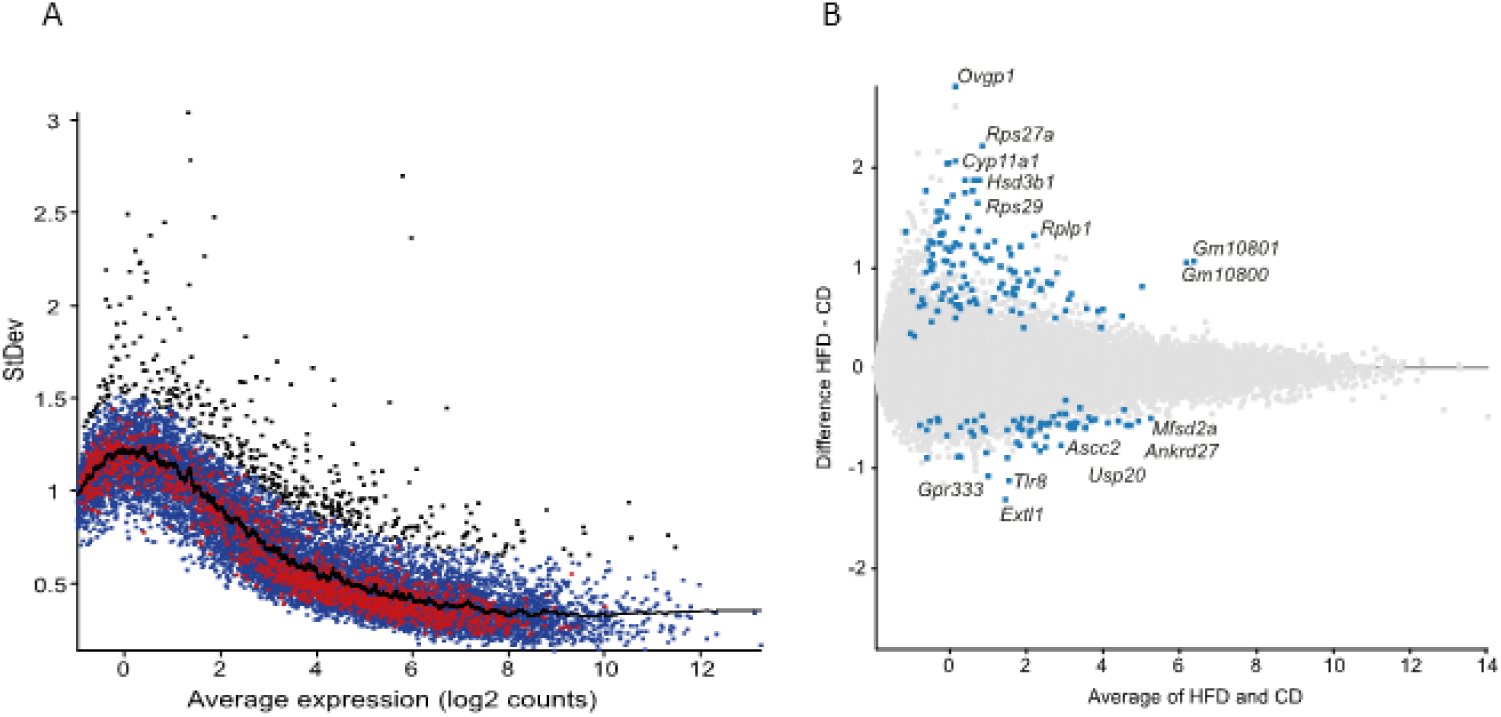
scRNA-Seq analysis of F1^mat-HFD^ oocytes. A. Variance plot highlighting in black the 198 most variable genes (> 0.528-fold from the mean standard deviation) in all samples. B. MA plot showing the difference in expression between HFD and CD of the 198 most variable genes (highlighted in blue).

## Methods

### Generation of targeted ESCs and mice

The *Dlk1-FLucLacZ* line was created by Taconic Biosciences and ESCs and animal founders were delivered to Imperial College.

### Maintenance of mice

Mice were handled and all *in vivo* studies were performed in accordance with the United Kingdom Animals (Scientific Procedures) Act (1986), were approved by the Imperial College AWERB committee and performed under a UK Home Office project license. Mice were housed on a 12-hour light-dark cycle with a temperature range of 21 +/-2°C in pathogen free conditions. The *Dlk1-FLucLacZ* line was maintained on a B6(Cg)-*Tyr*^*c-2J*^/J (C57Bl/6J albino) background. For mating, males were set up with not more than three females and morning plug checking was performed. Upon plug discovery, females were considered E0.5.

### Genotyping of animals

Genomic DNA was isolated from 4-week old ear biopsies or embryonic tails by digestion in lysis buffer (0.05 M Tris HCl pH 8, 0.025 M EDTA, 0.031% SDS, 0.02 M NaCl, 80 μg/ml Proteinase K (Sigma-Aldrich)) at 50°C with rocking. DNA was diluted 1:2 in 10 mM Tris HCl pH8 and 1 µl of diluted DNA was used in PCR analysis.

### Diet studies

Dlk1*-FLucLacZ* females were set up with B6(Cg)-*Tyr*^*c-2J*^/J males and upon vaginal plug discovery, matings were separated. Females were fed either a low protein chow (5769, TestDiet), a control chow (5755, TestDiet) or a high fat diet (56V6, TestDiet). All animals were returned to control diet at E18.5. Pregnant dams and embryos were imaged at E17.5 and offspring were imaged at P56. For multi-generational studies, *Dlk1-FLucLacZ* (males and females) mice that had been exposed to *in utero* high fat diet for the duration of pregnancy were aged to P56, with access to control diet *ad libitum*, imaged for bioluminescent activity and then set up with a B6(Cg)-*Tyr*^*c-2J*^/J partner. Offspring were aged to P56 prior to analysis.

### Bioluminescent imaging

D-Luciferin (Perkin Elmer) was dissolved in H_2_0 at 30 mg/ml. Mice were weighed and injected IP with 0.15 mg/g body weight, before being anaesthetized with isofluorane. Mice were imaged 10 mins post-injection, in an IVIS Spectrum (Perkin Elmer) under anaesthesia. Images of adult mice and pregnant dams were taken at field of view (FOV) C, with binning 4 and 180 sec exposure. For imaging of embryos, pregnant females were injected with D-Luciferin at least 12 mins prior to imaging. Embryos were dissected into 24 well dishes containing PBS and placed in the IVIS Spectrum. Images of embryos were taken at FOV A, with binning 4, focus 1 cm and 180 sec exposure. No additional D-Luciferin was added, and imaging continued for up to 35 mins post-injection. For *ex vivo* imaging of tissue, mice were culled at least 10 minutes after D-Luciferin injection, organs were removed and placed in clean dishes containing PBS. No further D-Luciferin was added to samples. Analysis of images was performed on Living Image software (Caliper Life Sciences). For quantification of bioluminescent signal, regions of interest were drawn around the whole abdomen and signal flux within the region was calculated.

### Beta-Galactosidase staining

E11.5 embryos were dissected and placed in cold LacZ fixative (2% formaldehyde, 0.2% glutaraldehyde, 0.02% Nonidet P-40, 1 mM MgCl_2_, 0.1 mg/ml Sodium Deoxycholate in PBS) for 4 hours, kept at 4°C with rocking. Tissue was washed in PBS before being placed in LacZ stain (0.4 mg/ml X-Gal, 4 mM Potassium Ferrocyanide, 4 mM Potassium Ferricyanide, 1 mM MgCl_2_, 0.02% Nonidet P-40 in PBS) for 4-6 hours at 4°C with rocking. Upon completion, embryos were washed twice in PBS before transfer to 70% ethanol and storage at 4°C. Photography was performed under standard light field conditions.

### Optical projection tomography

LacZ stained E11.5 embryos were mounted in cylinders of 2% low melting point agarose. The mounted samples were dehydrated through graded methanol solutions and maintained in 100% methanol prior to clearing. They were subsequently immersed overnight in an optical clearing solution, BABB (1:2 Benzyl benzoate: Benzyl alcohol, Sigma Aldrich). Optical projection tomography (OPT)^53^ was performed on a low-magnification imaging system as previously described^27^.

### RNA extraction and QRT-PCR analysis

RNA was extracted with TRIzol (Thermo Scientific) according to manufacturer’s protocol and all RNA precipitation steps were performed with 100% ethanol. Reverse transcription was performed using Superscript III Reverse transcriptase (Invitrogen) as per the manufacturer’s protocol, with minor modifications. RT-PCR was performed on a CFX96 Real-Time System (Bio-Rad) with QuantiTect SYBR Green Master Mix (Qiagen) as per the manufacturer’s protocol. Samples were normalised to *β-Actin* and expressed as the mean ± standard error.

### Preparation of MII-oocytes

Superovulation of 6-week old females was performed by evening IP injection of pregnant mare serum (PMS) followed by injection of human chorionic gonadotropin (HCG) 48hrs later. The following morning, mice were sacrificed via cervical dislocation and cumulus oophorus complex (COC) were collected from the oviduct via mechanical dissection. After digestion with hyaluronidase (Sigma Aldrich), MII oocytes were washed in sterile PBS and collected in RLT buffer (Qiagen). Oocytes were stored in 96-well plates at -80°C until further processed.

### Single-cell Bisulphite and RNA-Sequencing of MII-oocytes

Cell lysis was performed and Poly-A RNA was captured using oligo-dT conjugated to magnetic beads. Libraries were prepared and analysed according to the G&T-seq^54^ and Smart-seq2^55^ protocol. The lysate containing gDNA was purified on AMPureXP beads before bisulphite-sequencing (BS-seq) libraries were prepared and analysed according to the scBS-seq^36^ protocol that has previously been described.

### miR analysis of MII-oocytes

Total RNA was extracted from prepared oocytes with TRIzol (Thermo Scientific) according to manufacturer’s protocol. Small RNA library prep was performed with NEBNext Small RNA Library Prep (NEB) and microRNAs were isolated by size selection on 6% TBE PAGE gels 1mM (Novex) with clean up performed using Monarch PCR and DNA clean up kit (NEB), according to manufacturer’s protocol. Samples validation was performed on a Bioanalyzer 2100 using High Sensitivity DNA analysis kit (both Agilent) and sequenced on a MiSeq (Illumina).

### Bisulphite sequencing (tissue)

Bisulphite modification of DNA was carried out with the EZ Gold DNA Methylation Kit (Zymo Genetics) according to the manufacturer’s recommendations. PCR primers that specifically recognize bisulphite-converted DNA were used to amplify regions spanning the three imprinted DMRs associated with the *Dlk1-Dio3* imprinted region, while TaKaRa EpiTaq(tm) HS (Takara) was used for generic primers, using manufacturers protocol. PCR products were separated on an agarose gel and bands corresponding to the predicted size were excised and cleaned up with a Gel Extraction kit (QIAquick, Qiagen). Ligation of product was performed using Clone JET PCR cloning kit (Thermo Scientific) as per manufacturer’s protocol, before transformation into DH5α cells. Cells were plated onto LB/Ampicillin plates and grown up overnight at 37 °C. Colonies were picked (normally 24 per sample) and expanded in LB/Ampicillin broth overnight at 37°C. The following morning, plasmids were purified with the Wizard® SV 96 Plasmid DNA Purification System (Promega) according to the manufacturer’s protocol and sent for sequencing.

### Bisulphite primers

*Dlk1 sDMR* F: CCCCATCTAACTAATAACTTACA

R: GTGTTTAGTATTATTAGGTTGGTG

*IG-DMR* F: GTATGTGTATAGAGATATGTTTATATGGTA

R: GCTCCATAACAAAATAATACAACCCTTCC

*Gtl2 sDMR* F: GAAGAATTTTTTATTTGGTGAGTGG

R: CAACACTCAAATCACCCCCC

*PJET* R: CAGGAAACAGCTATGAC (sequencing)

### QRT-PCR Primers

*Dlk1 iso 1* F: GTACCCCTAACCCATGCGAG

*Dlk1 iso 2* F: TACCCCTAACCCATGCGAGA

*Dlk1 iso 3* F: TCCAGCACACCCAGGGAC

*Dlk1 iso 4* F: GCACACCCAGCCCGAG

*Dlk1 tg* R: GCCGGGCCTTTCTTTATGTT

*Dlk1 wt* R: CCCCGGTAATAGAGAAGGGC

*Dlk1 all F:* GAAAGGACTGCCAGCACAA

*Dlk1 all R:* CACAGAAGTTGCCTGAGAA

*B-Actin* F: CCTGTATGCCTCTGGTCGTA

*B-Actin* R: CCATCTCCTGCTCGAAGTCT

*Gtl2 F:* CGAGGACTTCACGCACAAC

*Gtl2* R: TTACAGTTGGAGGGTCCTGG

*Rtl1* F: TACTGCTCTTGGTGAGAGTGGACCC

*Rtl1* R: GGAGCCACTTCATGCCTAAGACGA

*Rtl1as* F: TCTCCACTCGAGGGTACTCCACCT

*Rtl1as* R: GTGGAGAACTTCGCTGTCATCGC

*Rian* F: ATGTCTGCTGCCCTGTCGTCT

*Rian* R: GCGGTCACTGCCAAGGTCTCT

*Mirg* F: GTTGTCTGTGATGAGTTCGC

*Mirg* R: GTTCTTGAACATCCGCTCC

### Genotyping primers

Dlk1 Geno F: AGTTTGCAAGCTGCACTTGG

Dlk1 Geno R: CTTTGGAGCTAGATCTTTCAGTGG

### Statistical Analysis

Student’s t-test was performed for statistical analysis of gene expression, microRNA analysis and flux calculations from *in vivo* imaging. For bisulphite sequencing analysis, Mann-Whitney tests were performed. For the analysis of variation among F2 animals, variance and 2-way ANOVA tests were performed.

### Author Contributions

MVDP, RMJ, AFS, MM, GK and AGF conceived and wrote up the study. MVDP, AG, SJM, WT, AD, CP, GM, LB, AS, ZW, JM, PF, AGU and JCF performed the mouse maintenance, experiments and analysis.

Materials and Correspondence: All requests should be made to Amanda G. Fisher

